# Whole Genome Assembly of a Hybrid *Trypanosoma cruzi* Strain Assembled with Nanopore Sequencing Alone

**DOI:** 10.1101/2023.07.27.550875

**Authors:** Jill M.C. Hakim, Sneider A Gutierrez Guarnizo, Edith Málaga Machaca, Robert H. Gilman, Monica R. Mugnier

**Affiliations:** Department of Molecular Microbiology and Immunology, Johns Hopkins Bloomberg School of Public Health, Baltimore, Maryland, USA; Department of International Health, Johns Hopkins Bloomberg School of Public Health, Baltimore, Maryland, USA; Asociacion Benefica PRISMA, Lima, Peru; Department of International Health, Johns Hopkins University Bloomberg School of Public Health, Baltimore, MD, USA

## Abstract

*Trypanosoma cruzi* is the causative agent of Chagas disease, which causes 10,000 deaths per year. Despite the high mortality caused by the pathogen, relatively few parasite genomes have been assembled to date; even some commonly used laboratory strains do not have publicly available genome assemblies. This is at least partially due to *T. cruzi’*s highly complex and highly repetitive genome: while describing the variation in genome content and structure is critical to better understanding *T. cruzi* biology and the mechanisms that underlie Chagas disease, the complexity of the genome defies investigation using traditional short read sequencing methods. Here, we have generated a high-quality whole genome assembly of the hybrid Tulahuen strain, a commercially available Type VI strain, using long read Nanopore sequencing without short read scaffolding. Using automated tools and manual curation for annotation, we report a genome with 25% repeat regions, 17% variable multigene family members, and 27% transposable elements. Notably, we find that regions with transposable elements are significantly enriched for surface proteins, and that on average surface proteins are closer to transposable elements compared to other coding regions. This finding supports a possible mechanism for diversification of surface proteins in which mobile genetic elements such as transposons facilitate recombination within the gene family. This work demonstrates the feasibility of nanopore sequencing to resolve complex regions of *T. cruzi* genomes, and with these resolved regions, provides support for a possible mechanism for genomic diversification.

## Introduction

*Trypanosoma cruzi* causes Chagas disease, a poorly understood and potentially fatal illness that is estimated to infect 6 million people worldwide. Chagas disease exhibits substantial phenotypic variability: only 30% of infected patients develop symptoms following chronic infection, which can involve different organ systems. Moreover, even parasite strains adapted to laboratory culture show phenotypic differences in drug susceptibility, *in vitro* growth capacity, and experimental infectivity in mice (Bice and Zeledon, 1970; Brener and Chiari, 1965; Rodriguez et al., 2014; Sykes et al., 2023). *T. cruzi’s* phenotypic diversity is accompanied by a corresponding genomic diversity that is observed across both field isolates and lab adapted strains (Llewellyn et al., 2011; Maiguashca Sánchez et al., 2020; Talavera-López et al., 2021; Wang et al., 2021). While it is not known what factors mediate *T. cruzi’s* phenotypic diversity, parasite genetic factors are thought to play a role (Macedo and Pena, 1998; Vago et al., 2000; Hakim et al., 2023).

Understanding which parasite genes contribute to disease and phenotypic variability first requires knowledge of *T. cruzi* genomic diversity. In the absence of full genome sequences, *T. cruzi* isolates are usually categorized into six genotypes termed Discrete Typing Units (DTUs), based on the results of several PCR and gel electrophoresis steps. However, these genotypes insufficiently describe the full gamut of parasite diversity and are not strongly associated with any clinical phenotype (Messenger et al., 2015; Zingales et al., 2012). Nevertheless, there are few high-quality whole genomes for *T. cruzi* available.

This is due, in part, to the fact that basic features of the parasite’s genome are difficult to study due to the genome’s repetitiveness, variability in genome sizes, number of chromosomes, spontaneous aneuploidies and polyploidies, and lack of synteny between strains. Moreover, two parasite genotypes (DTUs V and VI) are the results of hybridization between other parasite genotypes, resulting in highly heterozygous genomes that are even more difficult to resolve (Sturm et al., 2003). Additionally, what accounts for most of the genomic diversity between strains are 6 highly diverse and multicopy gene families, referred to as multi-gene families (MGFs) (Maiguashca Sánchez et al., 2020; Wang et al., 2021), which are distributed throughout the genome and thought to play important roles in immune evasion and parasite pathogenicity (Durante et al., 2017; Fonseca et al., 2019; Herreros-Cabello et al., 2020). These families further complicate genome assembly, as their multi-copy nature contributes to the repetitiveness of the genome, rendering them difficult to individually resolve or place within their genomic contexts.

Long read sequencing has increased in accuracy and accessibility in recent years. Recent *T. cruzi* genome assemblies have demonstrated the quality improvement of genomes assembled with long reads, but only when supplemented by additional Illumina sequencing (Berná et al., n.d.; Callejas-Hernández et al., 2018; Díaz-Viraqué et al., 2019; Wang et al., 2021). The long read approach allows the circumvention of several challenges posed by the *T. cruzi* genome: resolution of repetitive regions, accurate placement of diverse gene family members, as well as full resolution of transposable elements (TEs), which are difficult to study in most organisms. Like the multigene families, transposable elements are difficult to study using short reads alone: they are highly repetitive, can be multiple kilobases long, and can be distributed anywhere in the genome. *T. cruzi* has the highest proportion of transposable elements in its genome of any pathogenic kinetoplast, and transposable elements are hypothesized to play roles in both the unique transcriptional control of the parasite and in facilitating genomic rearrangements. (Bringaud et al., 2008, 2006; Gómez et al., 2021; Macías et al., 2018; Pita et al., 2019; Souza et al., 2007; Thomas et al., 2010).

We sought to develop a scalable pipeline for generating *T. cruzi* genomes to fully resolve these multigene families and TEs. We chose to develop this pipeline using the Tulahuen strain. The Tulahuen strain is commercially available from ATCC and has been used in numerous experimental studies, especially drug susceptibility studies due to its endogenously expressed LacZ reporter (Buckner et al., 1996; Mercado et al., 2019). Moreover, this parasite strain is a type VI DTU, and as such is one of the two hybrid DTUs, which are the most common genotypes in the Southern cone of the Americas, where the burden of Chagas disease is high. It is thus of great clinical relevance. Despite its relative importance, up to now there has been no publicly available genome for the Tulahuen strain, potentially limiting its experimental utility.

Here, we have used Oxford Nanopore Technology (ONT) long read sequencing, without supplementation with short Illumina reads, to generate a whole genome of the Tulahuen lacZ clone C4 strain. We find a high proportion of heterozygosity in the Tulahuen genome and annotate the genome for multigene family members and transposable elements. Using the newly annotated genome, we find that the distance between a transposable element and an open reading frame is bimodally distributed, and multi-gene family members fall exclusively within the histogram peak closest to TEs. This observation holds across multiple *T. cruzi* genomes, as well as in *Trypanosoma brucei*, suggesting that this genome organization is common to trypanosomes. This new genome provides critical information for basic biological research and raises interesting questions about the mechanism of virulence factor diversification.

## Methods

### Parasite culture

Trypanosoma cruzi epimastigotes (Tulahuen LacZ clone C4, BEI resources: NR-18959, DTU: VI) were grown in liver Infusion tryptose (LIT) medium supplemented with 10% heat-inactivated fetal bovine serum, 100 U/mL penicillin (Sigma-Aldrich, P3032), and 100 μg/mL streptomycin (Sigma-Aldrich, S9137). Parasites were seeded at a density of 1×106 cells/mL and harvested during the early stationary phase for DNA extraction.

Parasite DNA was extracted using the NEB HMW kit (NEB, T3050S), then left at 4C for at least two weeks prior to sequencing to relax tightly coiled DNA prior to sequencing. Using Ligation sequencing kits (112 and 114, catalog numbers SQK-LSK112, SQK-LSK114) ONT libraries were prepared and sequenced on two flow cells (9.4.1 and 10.4.1 respectively). The cumulative output from these two runs was 3.6Gbp and the N50 was 10kb. Raw Pod5 files were basecalled with Guppy super accuracy model, and duplex reads were called from data produced using the 10.4.1 flow cell, then integrated into the overall data.

### Genome sequencing and Assembly

A kmer histogram for 21-mers was generated with Jellyfish, then genome heterozygosity was estimated with GenomeScope (Marçais and Kingsford, 2011; Ranallo-Benavidez et al., 2020). Fastq data from 9.4.1 and 10.4.1 were pooled and were blasted against a database of known kDNA sequences. Raw reads that match kDNA were removed using seqkit, and the remaining reads were assembled using Nextdenovo, then polished using Nextpolish (Hu et al., 2023, 2020). Read coverage was assessed by mapping the raw reads to the assembly with minimap2 (Li, 2018). Busco was performed using the euglenozoan v10 database (Manni et al., 2021).

### Genome annotation

Coding sequences were annotated in two steps: First, genome annotations from the assembly of the Brazil A4 strain were mapped onto the new Tulahuen assembly using LiftOff (Shumate and Salzberg, 2021). Second, additional *de novo* gene calls were produced using AUGUSTUS trained on the Cl Brenner genome, which has fewer annotated genes than the Brazil strain, but is manually curated (Stanke and Morgenstern, 2005). These results were combined with GFFread -- merge function (Pertea and Pertea, 2020).

Multigene family members were further annotated using a method adapted from Wang et al (Wang et al., 2021). Briefly, a blast search against a database of known multi-gene family members was performed against the previously described coding regions with a percent identity cutoff of 85, and a length cutoff of 150 base pairs. If a genomic region matched multiple MGF members, the annotation was assumed to be non-specific and removed. To identify pseudogenes encoding for multigene family members, the search was performed against the entire assembly, and the genes that were found outside coding regions were defined as pseudogenes.

Transposable element annotation was performed using Extensive De Novo Transposon Annotator (EDTA) with the sensitive parameter (Ou et al., 2019).

All downstream analysis and figures were generated in R. All scripts used to generate the assembly, annotations, and analysis are available at https://github.com/mugnierlab/Hakim2023. Raw data and the final assembly will be deposited upon manuscript submission.

## Results

### Nanopore sequencing alone produces a complete whole genome

DNA from Tulahuen epimastigotes was extracted and sequenced using ONT R9.4.1 and R10.4.1 flow cells. Data from both runs were combined, and genome heterozygosity was estimated using GenomeScope. We describe a highly heterozygous genome, evident by a larger first peak in the kmer frequency distribution histogram, with an alternate allele frequency of 3.04% **(Fig1A)**. This high proportion of heterozygosity agrees with the ancestral history of this parasite strain: as a Type VI DTU, Tulahuen is a hybrid of two other parasite genetic types (cite). GenomeScope reports a haploid genome size of 48Mb, which is within the range of haploid genome sizes of other recently published genomes (45 and 53 MB; Supplemental table 1). After assessing heterozygosity, we assembled the genome using NextDenovo, resulting in a genome of 48.6 MB in 75 contigs, with an assembly N50 of 872KB and 57x mean coverage. Of the 75 contigs, 12 had telomeric repeats, and one contig had telomeric repeats at both ends, indicating one full chromosomal assembly.

**Figure 1:**
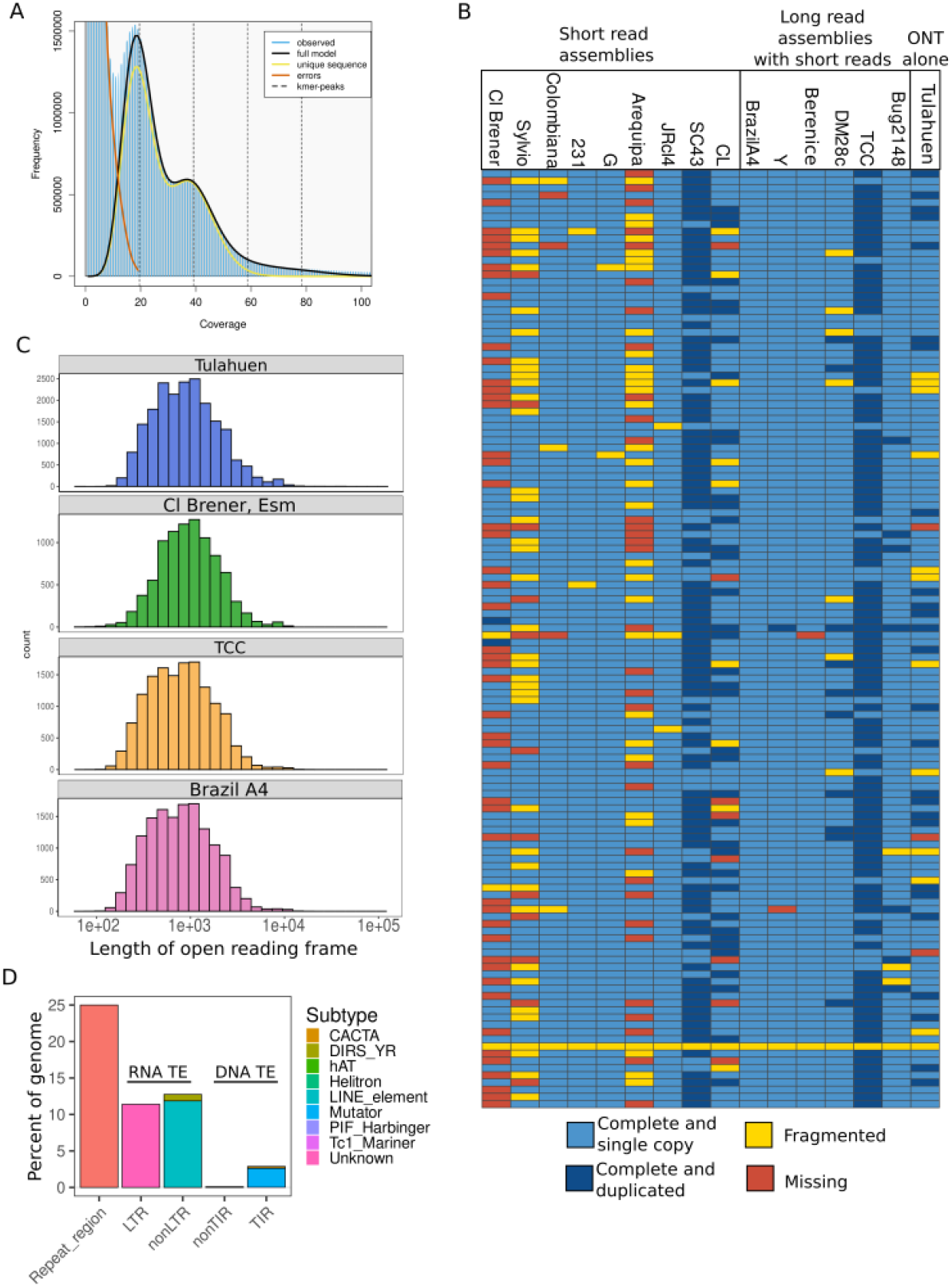
Whole genome assembly of a reference hybrid strain. A. Genomescope 21-mer histogram on raw nanopore reads. B. BUSCOs for newly assembled Tulahuen genome, as well as other *T. cruzi* genomes assembled with different data types. Columns are strains, rows are BUSCOs. C. Histogram of lengths for open reading frames in Cl Brener, Brazil A4, and Tulahuen genomes. D. Summary of TE classes found in the Tulahuen genome. CACTA = CACTA terminal inverted repeat, element, DIRS_YR = Tyrosine recombinase TE, LINE = Long Interspersed Nuclear Element, LTR = Long Terminal Repeat, TIR = Terminal Inverted Repeat

We then used BUSCO scores to assess assembly completeness based on the presence of conserved orthologous genes shared in the euglenozoan phylum. The Tulahuen assembly has a BUSCO completeness of 88.7%, which is slightly lower than recent assemblies using both short and long reads, though higher than most assemblies using short reads alone, including the reference CL Brener assembly (**Fig1B**). This suggests the need for updated assemblies using long read technologies. Notably, we find that one BUSCO gene, which encodes for UV excision repair RAD23-like protein, is reported as fragmented in every *T. cruzi* assembly assessed here, indicating that the gene is likely divergent from orthologs in other euglenozoan organisms.

It is important to note that BUSCO completeness, while a useful benchmarking tool for recovery of conserved, non-repetitive regions of the genome, may fail to accurately assess the resolution of more complex, repetitive regions of a genome, especially if the single copy genes are less likely to occur within repetitive regions. For example, our attempt at assembling this genome using Flye produced a genome that was 96% BUSCO complete, but only 26 MB long, suggesting that the conserved regions were well resolved in this assembly, but that much of the genome made up by repetitive regions, especially diverse MGF members and TEs, were lost. To more fully assess the accuracy of a *T. cruzi* genome assembly, additional analyses may be beneficial, especially ones tailored to a specific assembly method’s known systematic errors, such as ONT’s known issues in homopolymer resolution.

To evaluate systematic errors of the Nanopore only assembly, we took advantage of a NEO:LacZ construct cloned into the nuclear genome of the Tulahuen strain available from ATCC (Buckner et al., 1996). We found LacZ tandemly expressed ten times within the insertion locus: both the LacZ and Neo sequences share 99% identity with the insertion construct (3049/3050 and 790/792, respectively). However, all detected tandem LacZ and Neo constructs are identical to each other in this assembly, suggesting that the differences between the recovered sequences and the published constructs are likely true SNPs and not results of sequencing or assembly error. Importantly, we find no indels in this region, which is a known systematic error of Nanopore sequencing that typically requires short read supplementation to correct. Our ability to resolve this known sequence without indels is likely a result of the sequencing data from new 10.4.1 chemistry.

To further assess whether our assembly was affected by a systematic error that led to indels in the assembly, we compared the average open reading frame (ORF) length between this assembly and the reference genome, Cl Brener, and two other high quality long read assemblies, Brazil A4 and TCC. We chose Brazil A4 because of its high BUSCO completeness, and TCC because it is also a hybrid strain. Indels in low complexity regions are problematic when estimating ORFs, as indels will cause frame-shifts across the whole contig and result in erroneously short predicted open reading frames. We find that the distribution of ORF lengths is comparable for each genome **(Fig 1C)**.

### Transposable elements are physically distributed bimodally throughout the genome

Following assembly, we annotated the genome for repetitive regions that are generally difficult to resolve during assembly, specifically multi-gene family members and TEs. We find that a large proportion of the genome is made up of these elements, again in agreement with previous work: 25% are simple repeats, 27% are transposable elements, and 22.7% are MGF members. We found many RNA transposable elements, the most abundant of which were LINE and LTR retrotransposons **(Fig 1D)**. We also found DNA transposons, which have not previously been reported in *T. cruzi* genomes. However, the authors of TE annotation software used here have benchmarked their tool against high quality TE annotations, and found high false discovery rates for terminal inverted repeat (TIR) transposons (between 36 and 20% depending on the organism). Additional manual curation and experimentation will be required to confirm if these sequences are true DNA transposons.

To investigate the hypothesis that transposable elements contribute to diversification of multi-gene families, we evaluated the distribution of transposable elements in relation to open reading frames. If transposable elements were likely to act as donors for homologous recombination, facilitating the movement of multi-gene family members to different locations in the genome, we would expect them to be physically close to each other. We find that the distance between coding sequences and transposable elements is bimodally distributed, and that multi-gene family members are found exclusively in the first peak (**Fig2A**). This same pattern is observed in other assemblies, namely the high quality BrazilA4 (Pacbio and Hi-C) and Bernice (Nanopore and Illumina) genomes, indicating that this is unlikely to be an assembly artifact and rather represents a common aspect of trypanosome biology (S**up Figure 1**). Moreover, an additional analysis of *T. brucei* TEs and the highly variable, actively diversifying variant surface glycoprotein genes (VSGs) shows a similar pattern (**Fig2B**). Fewer VSGs make up the first peak for *T. brucei* compared to other ORFs; a closer look at the genes in this peak reveals many genes involved in TE biology, such as reverse transcriptase and RNAse H (**supplemental table 2**). This observation in a better annotated genome further supports the hypothesis that TE-mediated diversification may be evolutionarily conserved in trypanosomes.

**Figure 2:**
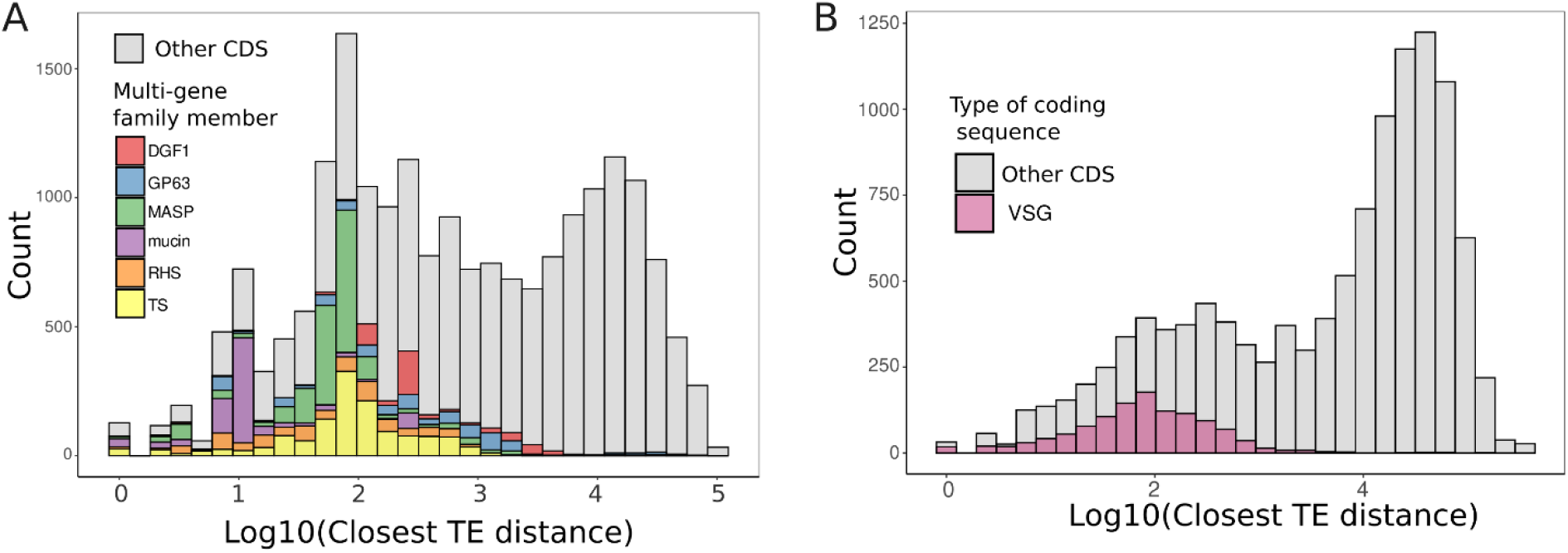
Transposable elements are associated with multigene family members. Histogram of the distance between the start or end of a transposable element to the start or end of any coding sequence (CDS) in the (A) Tulahuen T. cruzi genome and the (B) 427 T. brucei genome. In A, multi-gene family members are in stacked colored bars. All other CDSes are grey. In B, genes annotated as variable surface glycoproteins are in purple, and all other CDSes are in grey.

## Discussion

We have produced a full genome for a hybrid *T. cruzi* strain frequently used in biomedical research. Despite the complexity of this genome, it was assembled with relatively few reads: a total of 3.6Gbp, though 3.8% of those bases mapped to multi copy mitochondrial DNA, which was not used for the chromosomal assembly. This is the first *T. cruzi* genome that has been assembled with ONT long reads alone, without supplementation of other technologies, and the quality of this genome is comparable to those of other high-quality genomes. Low startup costs of Nanopore technology compared to Illumina or Pacbio, as well as the relatively low sequencing depth required, suggests that Nanopore sequencing may prove an excellent tool for generating whole genomes from a large number of strains, especially in low resource settings. The main systematic error, indels within homopolymers, seems to have a minimal effect on the predicted ORF lengths in this assembly; this is likely due to the new R10.4.1 sequencing technology, which in independent assessments shows improved resolution for these challenging regions (Sereika et al., 2022).

Using the long read assembly, we were able to annotate multi-gene family members and transposable elements and describe their relationship to each other in linear genome sequence. TEs in *T. cruzi* are known to be often clustered together(Olivares et al., 2000). Novel to this study is our observation that there seems to exist a genomic compartment containing coding regions that are isolated from TEs; this could point to negative selection keeping potentially deleterious insertions and rearrangements away from important and conserved genome compartments.

Unlike other highly variable gene families, such as VSGs in *T. brucei* and var genes in *P. falciparum*, multi-gene families in *T. cruzi* are not primarily localized to the subtelomere, which is often a hotspot for diversification driven by mitotic recombination. The dispersed nature of *T. cruzi*’s variable genes suggests an alternative, or additional, mechanism for diversification. The data presented here and elsewhere suggest a possible role for TEs in this putative mechanism. There is other evidence of TE’s association with MGFs: TcTREZO, a site-specific retrotransposon, shows frequent insertion into MASP genes, and is generally found in non-syntenic regions of the *T. cruzi* genome containing MGFs (Souza et al., 2007). Further genomic analysis of actively diversifying strains isolated from field samples is required to fully characterize the diversity within these multi-gene families, and additional experimental strategies will help uncover the specific mechanisms underlying any TE-mediated diversification processes.

Very few whole genomes of *T. cruzi* exist, despite the parasite’s substantial genetic diversity and the putative role parasite genetics may play in disease heterogeneity. Further, even fewer genomes of hybrid *T. cruzi* genomes exist: these strains are more genomically complex, but critical to understanding parasite diversity, as the majority of parasites in the southern cone of the Americas are hybrid strains. This work demonstrates the feasibility of using Nanopore sequencing alone, with relatively little sequencing data to study important genomic features such as multi-gene family members and transposable elements in a hybrid strain. These new and improving tools will allow us to generate more whole genome data for this enigmatic and genetically unstable parasite.

## Acknowledgements

JMCH is supported by T32AI007417 and a JHU Discovery Award. SAGG is supported by R01AI151295. RHG is supported by R01AI107028 and R01AI151295. EMM is supported by D43 TW001140. MRM is supported by R01AI151295 and a JHU Discovery award.

## Notes

https://github.com/mugnierlab/Hakim2023

### Competing Interest Statement

The authors have declared no competing interest.

